# Hippocampal Volume, Neurite Density and Trace Eye-Blink Conditioning in Young Children

**DOI:** 10.1101/2022.01.03.474711

**Authors:** Vanessa Vieites, Yvonne Ralph, Bethany Reeb-Sutherland, Anthony Steven Dick, Aaron T. Mattfeld, Shannon M. Pruden

## Abstract

The current study examined the relations between hippocampal structure (e.g., volume and neurite density) and performance on a trace eye blink conditioning (EBC) task in young children. Our first aim assessed whether individual differences in hippocampal volume were associated with trace EBC performance, using both percent Conditioned Responses (% CR) and CR onset latency or the average latency (ms) at which the child started their blink, as measures of hippocampal-dependent associative learning. Our second aim evaluated whether individual differences in hippocampal neurite density were associated with EBC performance using the same outcome measures. Typically developing 4- to 6-year-olds (*N* = 31; 14 girls; *M_age_* = 5.67; S_Dage_ = 0.89) completed T1 and diffusion-weighted MRI scans and a 15-minute trace eyeblink conditioning task outside of the scanner. % CR and CR onset latency were computed across all tone-puff and tone-alone trials. While hippocampal volume was not associated with any of our EBC measures, greater hippocampal neurite density bilaterally, was associated with later CR onset. In other words, children with greater left and right hippocampal neurite density blinked closer to the US (i.e., air puff) than children with less hippocampal neurite density, indicating that structural changes in the hippocampus assisted in the accurate timing of conditioned responses.

## Introduction

The hippocampus, a medial temporal lobe structure, plays a vital role in memory (Squire et al., 2004). Human hippocampal synaptic connectivity matures around five years of age (Seress, 2001) with structural magnetic resonance imaging (MRI) indicating development into early adulthood (Gogtay et al., 2006). Protracted development is associated with later onset of hippocampal-dependent behaviors (Jabès & Nelson, 201). Yet, in early childhood the relation between hippocampal development and memory is understudied (Bauer et al., 2017).

Functional magnetic resonance imaging (fMRI) facilitates the investigation of brain and behavior relations, particularly for subcortical regions (e.g., hippocampus and memory development), but is sensitive to movement artifact. Young children have difficulty remaining motionless while engaged in tasks, making it difficult to obtain valid fMRI data. Structural MRI can be used with young children and provides data on hippocampal volume using high-resolution T1- and T2-weighted structural MRI and neurite density using neurite orientation dispersion and density diffusion weighted imaging (NODDI-DWI). While structural MRI is a useful tool for studying the morphology of neural regions, it does not directly measure hippocampal function, which raises challenges for exploring how hippocampal structure relates to behavior. The present study addresses this challenge by collecting structural MRI data along with behavioral data outside of the scanner in young children using a developmentally appropriate task, trace eyeblink conditioning (EBC), that assesses the efficiency of learning processes supported by the hippocampus (Kim et al., 1995; Büchel et al., 1999).

EBC entails learning associations between auditory and tactile stimuli (Herbert et al., 2003). During EBC a conditioned stimulus (CS; tone) is paired with an unconditioned stimulus (US; eye air puff) that elicits a reflexive blink or an unconditioned response (UR). Following several pairings, the tone CS alone elicits a blink or conditioned response (CR). In trace EBC, the offset of the CS and onset of the US are separated by a “trace” period (Büchel et al., 1999). Trace EBC is hippocampal-dependent and may serve as a proxy for hippocampal functioning in very young children (Vieites et al., 2020). Hippocampal lesions result in poorer learning during trace EBC (Kim et al., 1995) and fMRI studies reveal significant hippocampal activation during trace EBC conditioning (Büchel et al., 1999).

The current study examined the relations between hippocampal structure (e.g., volume and neurite density) and performance on a trace EBC task in young children. Our first aim assessed whether individual differences in hippocampal volume were associated with trace EBC performance, using both *percent CRs* and *CR onset latency* as measures of hippocampal-dependent associative learning. Our second aim evaluated whether individual differences in hippocampal neurite density were associated with EBC performance using the same outcome measures.

## Method

Typically developing 4- to 6-year-olds (*N* = 31; 14 girls; *M_age_* = 5.67; *S_Dage_* = 0.89; 42% Hispanic, 13% African American or Black, 6% White non-Hispanic, and 39% mixed race or another ethnicity) completed T1 and diffusion-weighted MRI scans and a 15-minute trace eyeblink conditioning (EBC) task outside of the scanner (see detailed methods for additional information). Children first participated in a practice scan in a realistic “mock” scanner. Once children were acclimated, they completed the 30-minute MRI scan. Two-dimensional surface renderings of each participant’s T1-scans were constructed using Freesurfer v6.0 (Dale et al., 1999). Regions of interest, including the hippocampus and cerebellum, were defined anatomically using the semiautomated Freesurfer parcellation procedure (Fischl et al., 2004). An estimate of intracranial volume was also obtained (Buckner et al., 2004). In the present study we also focused on the neurite density index (NDI) in grey matter, higher values of NDI correlate with higher density of neuronal tissue (Shao et al., 2021). We computed the NDI using the Microstructure Diffusion Toolbox (Harms et al., 2017). Using the Freesurfer ROI segmentation as a mask, volume and NDI were calculated for the hippocampus and cerebellar cortex bilaterally.

Following the MRI scan, children participated in a 15-min trace EBC task that consisted of 80 tone-puff trials (i.e., presentation of a 750-ms, 80dB, 1000 Hz pure tone CS, followed by a 500-ms interstimulus trace interval during which no stimulus was presented, followed by a 100- ms air puff US, ~10 lb/in^2^), 10 tone-alone trials, and 10 puff-alone trials (Figure 1; Jacobson et al., 2011). Two trace EBC associative learning variables were calculated. Conditioned responses (*% CRs*) were defined as eyeblinks that occurred between 800 and 1300 ms after the onset of the tone during the tone-puff paired trials and after 800 ms during the tone-alone trials. *CR onset latency*, or the average latency (ms) at which the child started their blink, was computed across all tone-puff and tone-alone trials.

**Figure 1.**
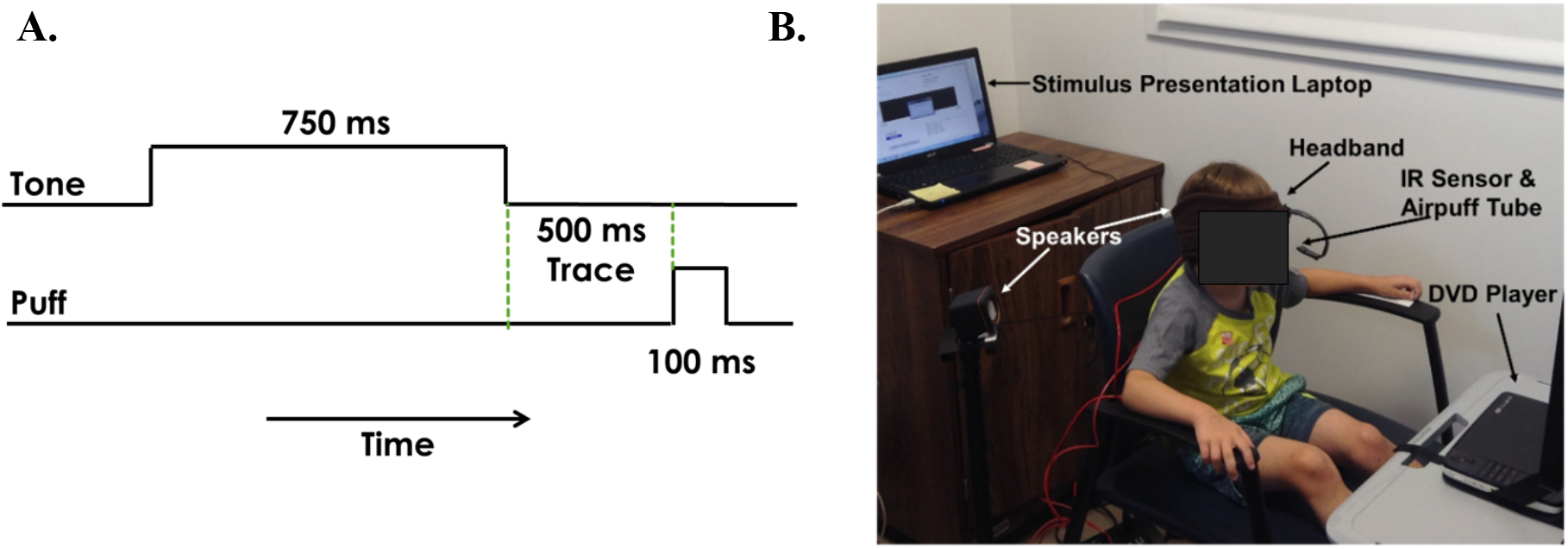
Trace EBC paradigm depicting times of CS, trace period, and US (A) and EBC setup (B). Panel B shows a young child facing (face blinded for anonymity) a DVD player while wearing a headband with a sensor and tube pointing toward their left eye. The tube delivers the air puffs and the sensor measures blink responses. One speaker is positioned on each side of the child’s head approximately 12 inches away.

## Results

We first examined whether hippocampal volume was related to % *CRs* and *CR onset latency*, controlling for child age, birth sex, maternal education, and intracranial volume. Robust linear regression analyses (using a Huber loss function implemented in R package *rlm*) showed that hippocampal volume was not associated with *CR onset latency* (Left: *p* = 0.61; Right: *p* = 0.70), or % *CRs* across all tone-puff paired and tone-alone trials (i.e., 90 trials; Left: *p* = 0.83; Right: *p* = 0.72) or just tone-alone trials (i.e., 10 trials; Left: *p* = 0.61; Right: *p* = 0.19).

In contrast to volume, robust linear regressions with hippocampal NDI, controlling for child age, birth sex, maternal education, and cortical NDI, showed greater neurite density of the left (*B* = 1724.41 [95% CI 832.14 to 2531.10]; *t*(24) = 3.88, *p* = 0.0007, semipartial *r* (*r_sp_* = 0.52) and right (*B* = 1420.11 [95% CI 484.28 to 2355.94]; *t*(24) = 2.97, *p* = 0.007, *r_sp_* = 0.47) hippocampus was associated with higher *CR onset latency* (Figure 2, top). However, bilateral hippocampal neurite density was not associated with % *CRs* across all 90 paired and tone-alone trials (Left: *p* = 0.13; Right: *p* = 0.17) or across the 10 tone-alone trials (Left: *p* = 0.76; Right: *p* = 0.78).

**Figure 2.**
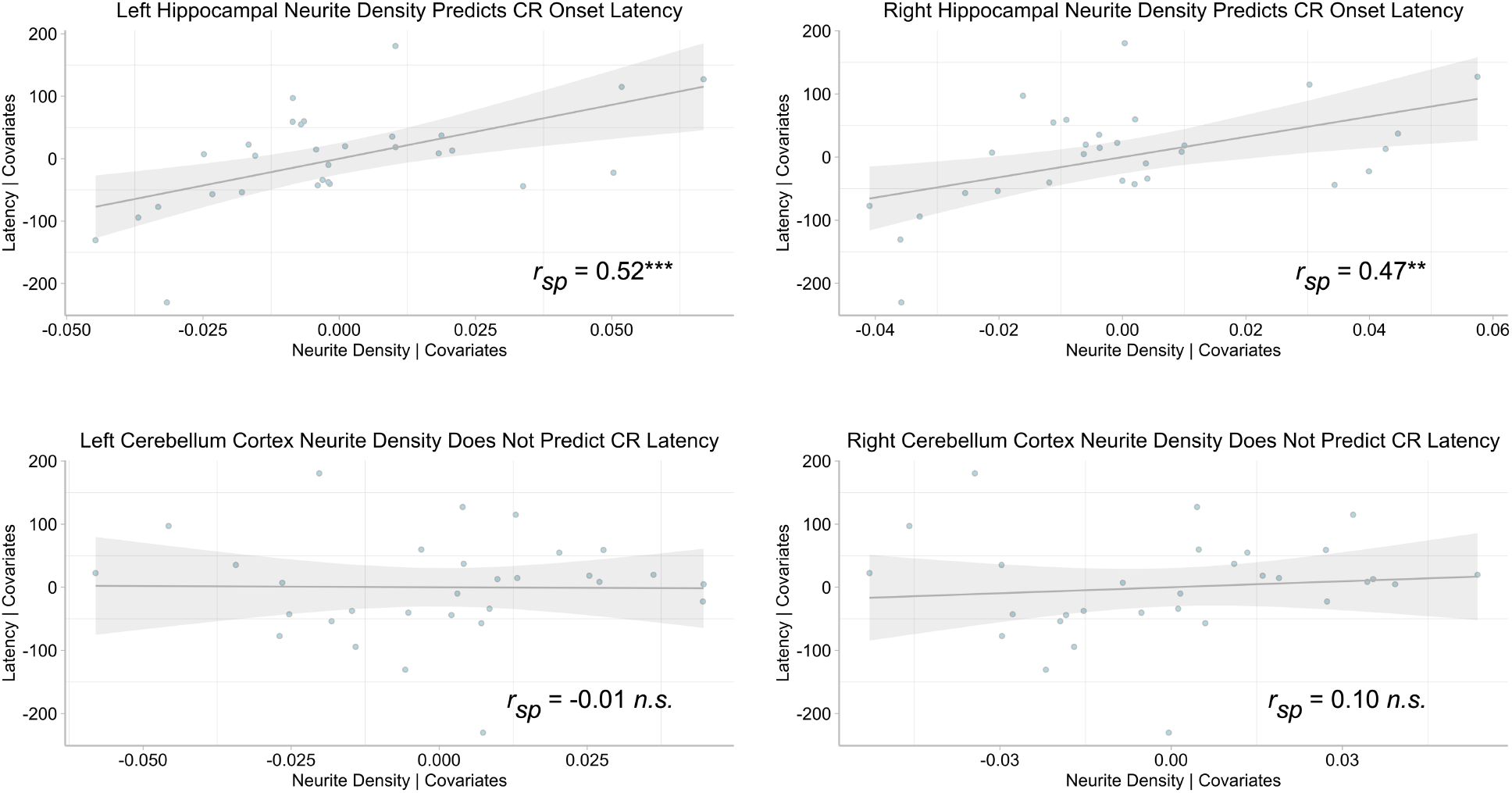
Robust linear regression added-variable plots show the association between *CR onset latency* and left and right hippocampal neurite density index (*NDI*; top) and left and right cerebellar cortical NDI (bottom), controlling for child age, birth sex, maternal education, and cerebral cortical NDI covariates. Shaded area shows the 95% Confidence Interval (*CI). r_sp_* = semipartial correlation showing the correlation between NDI and conditioned response latency controlling for variance associated with covariates. *n.s*. = non-significant; \*\**p* < .01; *** *p* < .001.

To ensure our findings were unique to the hippocampus and not to other subcortical regions potentially involved with trace EBC, we also examined cerebellar NDI in relation to *CR onset latency*. The cerebellum has been implicated in delay (Thompson & Krupa, 1994) and trace (Woodruff-Pak et al., 1985) EBC. Neither left (*B* = 119.42 [95% CI −958.63 to 1197.47]; *t*(24) = 0.22, *p* = 0.83, semipartial *r (r_sp_* = −0.01) nor right (*B* = 523.94 [95% CI −377.29 to 1425.16]; *t*(24) = 1.14, *p* = 0.27, *r_sp_* = 0.10) cerebellar cortical ND, controlling for child age, birth sex, maternal education, and cortical NDI, were associated with *CR onset latency*.

## Discussion

Considering the challenges in obtaining neural correlates of behavior in young children, the current study sought to assess the validity of using a hippocampal-dependent trace EBC paradigm as a proxy for hippocampal function by investigating the neural correlates of both trace EBC performance (% CRs) and CR onset latency. This study bridges our understanding of hippocampal development and behavioral performance (Vieites et al., 2020) and is among a few to investigate trace EBC as a viable proxy for hippocampal function in young children (Vieites et al., 2015).

While hippocampal volume was not associated with any of our EBC measures, greater hippocampal neurite density bilaterally, was associated with *later CR onset*. In other words, children with greater left and right hippocampal neurite density blinked closer to the US (i.e., air puff) than children with less hippocampal neurite density, indicating that structural changes in the hippocampus, assessed by NDI, assisted in the accurate timing of conditioned responses. This finding aligns with previous work with adults using fMRI and MEG which indicate that the hippocampus is activated during trace EBC associative learning (Büchel et al., 1999; Kirsch et al., 2003) and that it may be specifically responsible for accurate timing of the conditioned response (Knight, 2004). The timing of eyeblinks is critical to normal acquisition of learned associations (Boneau, 1958), the purpose being to blink just prior to the onset of the US so that the eye is protected from the air puff.

The current study is not without limitations. Our data were correlational and thus, do not allow us to infer the direction of the relation between hippocampal development and function. In addition, a larger sample size may have yielded significant results that were not present (e.g., between % CRs and hippocampal volume or NDI). Despite these limitations, this study provides an important first step toward better understanding the relations between hippocampal development and hippocampal-dependent cognitive abilities in early childhood. Participants with a greater left and right hippocampal NDI exhibited more accurate timing in their CRs. These results suggest that hippocampal maturation, captured by changes in neurite density may engender resource-efficient differentiation of signal from noise in learned associations. As such, trace EBC may be a novel non-invasive and simple proxy for investigating hippocampal function in young children.

### Detailed Methods

Imaging was performed using a 3-T Siemens MAGNETOM Prisma MRI scanner (V11C) with a 32-channel coil. We collected structural anatomical scans using a whole-head 3D T1-weighted acquisition inversion prepared RF-spoiled gradient echo protocol (93 axial slices at 1 mm isotropic resolution) with prospective motion correction (Siemens vNAV; Tisdall et al., 2012). We also collected multi-shell high-angular diffusion-weighted imaging (HARDI) data according to the Adolescent Brain and Cognitive Development (ABCD) protocol (Hagler Jr et al., 2019). These scans were collected with a 1.7 mm isotropic voxel size, using multiband imaging echo planar imaging (EPI; acceleration factor = 3). The acquisition consisted of ninety-six diffusion directions, six b=0 frames, and four b-values (102 diffusion directions; 6 b=500 s/mm^2^, 15 b=1000 s/mm^2^, 15 b=2000 s/mm^2^, and 60 b=3000 s/mm^2^).

Initial post-processing of the diffusion data was accomplished with DTIPrep v1.2.8 (Oguz et al., 2014), TORTOISE DIFFPREP v3.1.0 (Irfanoglu et al., 2017; Pierpaoli et al., 2010), AFNI (v 20.6.02), and FSL v6.0.1 topup (Andersson et al., 2003). We also implemented a pre- and post-analysis quality check assessing signal-to-noise of each diffusion b-value (Roalf et al., 2016). Initial quality control was accomplished in DTIPrep to complete the following steps: 1) image/diffusion information check; 2) padding/cropping of data; 3) Rician noise removal; 4) slicewise, interlace-wise, and gradient-wise intensity and motion checking. TORTOISE DIFFPREP was used to accomplish motion and eddy current correction, and registration to the T1-weighted structural scan, which was maintained in original subject space.

NODDI is an alternative diffusion model that can distinguish among three tissue-property contributions to the diffusion signal: intracellular, extracellular, and cerebrospinal fluid (Zhang et al., 2012).With respect to the present study, the NODDI model allows estimation of the contributions of neurite morphology from the diffusion signal, and such estimates such as neurite density from the NODDI model have been verified with histology in animals (Sato et al., 2017) and pathological findings in humans (Sone et al., 2020).

Trace EBC stimulus presentation, data collection, and data processing were completed using a commercially available human eyeblink conditioning system (San Diego Instruments, San Diego, CA). Children were fitted with a soft headband, attached to an infrared emitter-sensor that recorded their eyeblink responses and tubing that delivered the air puffs. The sensor and tubing were positioned approximately 2 inches from the child’s left eye. Tones were delivered through external speakers positioned 12 inches away from each side of the child’s head.

